# Gene expression profiling for forensic age assessment of porcine skin wounds

**DOI:** 10.1101/2025.11.06.686914

**Authors:** Cecilie Bækgård, Kerstin Skovgaard, Marie Høy Hansen, Henrik Elvang Jensen, Kristiane Barington

## Abstract

Determining the age of wounds is of utmost importance in both veterinary and human forensic pathology. The aim of this study was to design and optimize quantitative polymerase chain reaction (qPCR) primers for use on degraded RNA samples obtained in veterinary forensic cases, hereby ensuring assay robustness. Moreover, the aim was to evaluate if an expression signature, based on optimized short-amplicon primers, was able to differentiate porcine experimental granulation tissue according to age and if this could be used for wound age assessment in veterinary forensic cases.

Initially, 12 samples of experimental granulation tissue (n=6) and skin (n=6) from two pigs were deliberately exposed to RNA degrading conditions before being stored in RNAlater. A panel of 24 robust primers were selected based on the intentionally degraded samples.

Granulation tissue (5, 10, 15, 20, 25, 30, and 35 days of age) (n=94) and control skin (n=47) from 47 experimental pigs was sampled and stored in RNAlater. Furthermore, granulation tissue and fibrous scar tissue were sampled from 14 veterinary forensic cases. Microfluidic qPCR was performed to evaluate the gene expression of 24 genes. An expression signature of 14 genes reflected the age of the experimental wounds. The 5-day old wounds displayed the biggest divergence from the control skin. As the granulation tissue matured, gene expression gradually approached the levels observed in intact skin. The forensic samples clustered somewhat separately from the experimental samples.

In conclusion, granulation tissue from the experimental wounds displayed a time-dependent expression profile based on 14 short-amplicon primers suitable for use with low-quality RNA. However, an expression profile of 14 genes cannot be used as the sole method for forensic age assessments of porcine wounds.

## Introduction

Veterinary pathologists are commonly required to carry out forensic examinations of animals with skin wounds. In these cases, wound age can reflect the degree of neglect or be useful to determine in whose custody the animal was when the lesion was inflicted. Therefore, determining the precise age of wounds is of utmost importance (Barington and Jensen, 2013, Knight and Saukko, 2016).

In Denmark, approximately 80% of veterinary forensic cases concern pigs and in approximately one third of these cases wounds are the primary lesion (Barington, 2018, Pankoke et al., 2023). The majority of these wounds are caused by external trauma and though some are only a few hours old, most are chronic and characterized by grossly visible granulation tissue. Presently, the presence and thickness of the granulation tissue formed during healing is used to estimate the age of these wounds (Barington et al., 2016a, Barington et al., 2016b). Barington et al (2018) observed an increase in granulation tissue thickness up to day 10, after which thickness decreased again (Barington et al., 2018). This study showed that granulation tissue thickness is not linearly correlated with wound age. Age estimations in veterinary forensic cases are often given as “several days”, “several weeks” or “several months” (Barington et al., 2018).

Despite the numerous methods and techniques, including, but not limited to, histology, immunohistochemistry and quantitative polymerase chain reaction (qPCR), that have been employed to study age assessment of skin wounds it continues to be a challenge obtaining an accurate age estimates of wounds (Barington et al., 2016a, Barington et al., 2018, Barington et al., 2017, de Siqueira et al., 2016, Li et al., 2020, Pankoke et al., 2023). The most common models to study wound healing are murine models. These models are often chosen due to their low-cost, easy handling and the wide selection of readily available reagents (Masson-Meyers et al., 2020, Saeed and Martins-Green, 2023). In addition, numerous tools are available for the genetic manipulation of the mouse model allowing for the study of specific aspects of wound healing (Beare et al., 2003, Short et al., 2022, Tomasek et al., 2013, Toriseva et al., 2012). Mice and rats, however, heal mainly by contraction, whereas humans and pigs heal mainly by re-epithelialization (Boyko et al., 2017). Due to the anatomical and physiological similarities of human and porcine skin, it is likely that a porcine wound healing model is useful not only in the age prediction of porcine wounds, but also comparable to wounds in humans (Parnell and Volk, 2019, Saeed and Martins-Green, 2023, Sullivan et al., 2001). Porcine models have previously been used to study wound healing. However, these models have focused on wound treatments (e.g. topical treatments, bandages) and not time dependent changes in wounds healing per secundam (Baetz et al., 2023, Dearman et al., 2023, Patil et al., 2018, Petersen et al., 2016, Plettig et al., 2015). As a result, many experimental porcine wound models do not accurately reflect the types of wounds observed in veterinary forensic cases.

Microfluidic qPCR is a powerful technique for quantifying gene expression levels in nanolitre volumes across a large number of samples simultaneously, allowing for the identification of molecular markers associated with different stages of granulation tissue development. In recent studies, mRNA expression profiles have been created from an experimental porcine bruise model in which Barington et al discovered time-dependent expression patterns in porcine bruises (Barington et al., 2017). However, the study did not account for the fact that samples taken from forensic cases often are of poor RNA quality. Therefore, these gene expression profiles could not be used for age assessment in a substantial part of forensic cases concerning bruises (29).

In order to develop a gene expression signature applicable in forensic wound age assessment an assay based on short-amplicon primers suitable for use with low-quality RNA, targeting time dependent mRNA markers must be designed and optimised. In the present study, transcripts known to play roles in inflammation, extracellular matrix remodelling, angiogenesis, cellular proliferation and wound closure were selected for the above mentioned gene expression signature. The primary aim of the study was to design short-amplicon primers suitable for use on degraded RNA samples like those of veterinary forensic cases. A second aim of the study was to evaluate if the expression signature based on selected genes were able to differentiate porcine experimental granulation tissue according to age. Moreover, to investigate if such a gene expression signature could be used to determine the age of granulation tissue in veterinary forensic cases.

## Materials and methods

### Animal model

A total of 50 female specific pathogen free Yorkshire-Landrace crossbred pigs were used in this study. Each of the pigs had a bodyweight of 20-25 kg and were allocated into seven groups in the order they arrived at the facility. Three pigs were excluded from the study, two of which died during anaesthesia, and one was euthanized within 12 hours after surgery due to respiratory distress.

The experimental procedure was carried out as previously described (Bækgård et al., 2025). In summary, full thickness wounds were created by incision on the back of each pig (Figure 1). All wounds were left to heal by second intention for 5, 10, 15, 20, 25, 30, or 35 days (Table 1). Skin excised from wound area A was sampled as control tissue and stored in RNAlater at 5°C for 24h before being stored at -20°C until RNA extraction.

**Fig. 1:**
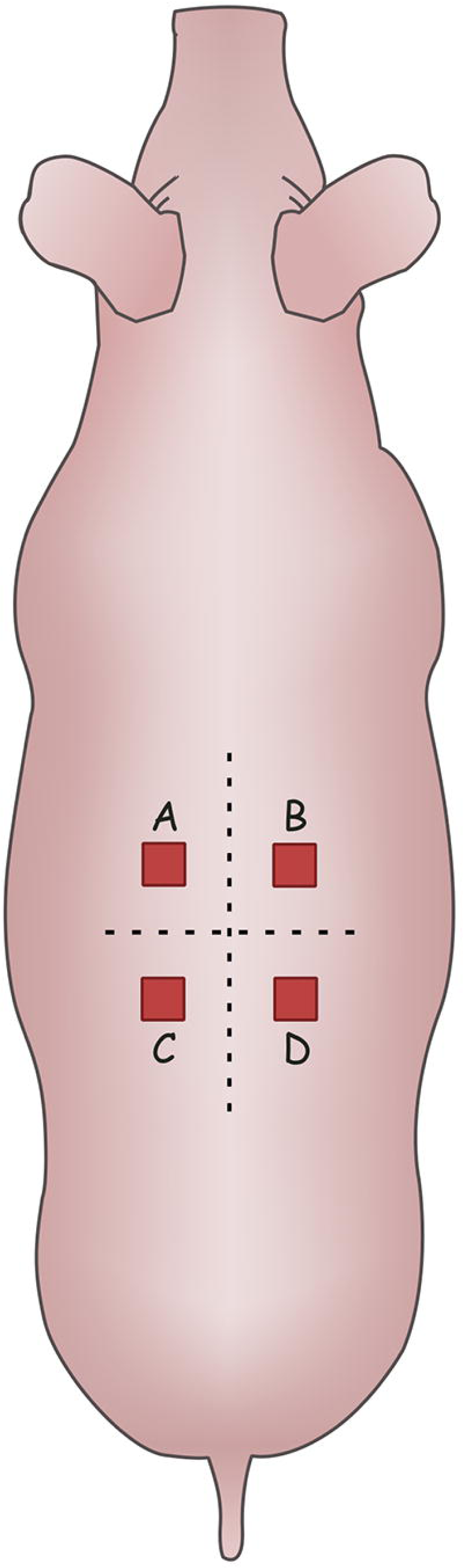
Localization of wounds A-D. The wounds were placed 4 cm lateral to the spine and 4 cm cranial or caudal to the last rib. Skin was sampled as control from wound A. Samples of granulation tissue from wounds A and D were collected postmortem. Figure made using Inkscape.

**Table 1.** Degraded sample overview. Sample type, number of samples and treatment of samples prior to being stored in RNAlater at 5°C for 24h before being stored at -20°C until RNA extraction.

| Sample type | No. of samples | Treatment |
| --- | --- | --- |
| Skin | 2 | Frozen, then thawed |
| Skin | 2 | Room temperature for 24 hours |
| Skin | 2 | Room temperature for 48 hours |
| Granulation tissue | 2 | Frozen, then thawed |
| Granulation tissue | 2 | Room temperature for 24 hours |
| Granulation tissue | 2 | Room temperature for 48 hours |

The pigs were euthanized at different timepoints (5, 10, 15, 20, 25, 30, or 35 days post wounding). Postmortem, wounds A and D were cross sectioned and granulation tissue was samples from the center of the wounds before being stored in RNA later under the same conditions as the control skin. Additionally, the thickness of the granulation tissue was measured to be from 0.3 to 1.2 cm (*x̅*= 0.6 cm) (Bækgård et al., 2025).

### Degraded samples

An additional 12 samples of skin and granulation tissue were sampled and treated to purposely degrade the RNA before being stored in RNAlater (Table 1). These samples consisted of excised skin from surgery site A and granulation tissue from wounds B and C from two pigs. Three methods were applied to intentionally degrade the samples: they were frozen and then thawed or left on a lab bench at room temperature for either 24 or 48 hours. Subsequently, all samples were stored in RNAlater at 5°C for 24h before being stored at -20°C until RNA extraction.

### Forensic cases

In addition to the experimental wounds, 28 forensic wounds were sampled (Table 2). Fourteen cases of porcine skin wounds were received for forensic examination at the University of Copenhagen and processed from January to September 2021. All fourteen cases were reported by the Danish Veterinary and Food Administration or the Danish Agriculture Agency. Samples were taken from the center of the wounds and divided in granulation tissue and underlying fibrous scar tissue whenever possible before being stored in RNAlater in the same manner as for the experimental samples. Information regarding estimated wound age, locations and sizes of the wounds, and thickness of granulation tissue and fibrous scar tissue were derived from forensic reports.

**Table 2.** Overview of veterinary forensic samples. A total of 28 forensic samples from 14 different cases originating from various locations on the pigs were included in the study. The wound age given is the one stated in the forensic report. GT: granulation tissue, FST: fibrous scar tissue, RIN: RNA Integrity Number. *Total thickness of GT and FST.

| Case | Wound number | Wound location | Slaughter process/<br>frozen | Max. length (cm) | Wound age | No. of samples collected | Type of sample (GT/ FST) | Thickness of GT / FST (cm) | RIN |
| --- | --- | --- | --- | --- | --- | --- | --- | --- | --- |
| 1 | 1 | Snout ridge | Yes/yes | 14 | 4-5 days | 1 | GT | 0.2 / - | 3.2 |
| 2 | 2 | Abdomen (umbilical hernia) | No/yes | 5.3 | > weeks | 2 | GT + FST | 0.1 / 0.4 | 2.4<sup>GT</sup><br>2.4<sup>FST</sup> |
| 3 | 3 | Abdomen (umbilical hernia) | Yes/yes | 14 | > weeks | 2 | GT+ FST | 0.2 / 3.5 | 3.3<sup>GT</sup><br>4.2<sup>FST</sup> |
| 4 | 4 | Abdomen (umbilical hernia) | Yes/yes | 9 | > weeks | 2 | GT + FST | 0.3 / 1 | 2.7<sup>GT</sup><br>5.6<sup>FST</sup> |
| 5 | 5 | Abdomen (umbilical hernia) | Yes/yes | 12,5 | > weeks | 2 | GT + FST | 0.9 / 1 | 6.7<sup>GT</sup><br>5.8<sup>FST</sup> |
| 6 | 6 | Abdomen (herniating enterocytoma) | Yes/yes | 7.5 | > 1 week | 2 | GT + FST | 0.4 / 0.3 | 5.7<sup>GT</sup><br>2.7<sup>FST</sup> |
| 7 | 7 | Tail | Yes/yes | 7 | > 1 week | 1 | GT | 0.4 / - | 2.2 |
| 8 | 8 | Tail | Yes/yes | 2.1 | > weeks | 1 | GT | 0.4-0.6 / - | 2.7 |
| 9 | 9 | Tail | Yes/yes | 3.6 | > weeks | 1 | GT | 1 / - | 2.8 |
| 10 | 10 | Tail | No/yes | 3.5 | > weeks | 2 | GT + FST | 1* | 2.6<sup>GT</sup> |
|  |  |  |  |  |  |  |  |  | 5.3<sup>FST</sup> |
| 11 | 11 | Tail | No/yes | 3 | > weeks | 2 | GT+ FST | 0.5 / 1 | 2.6<sup>GT</sup> |
|  |  |  |  |  |  |  |  |  | 2.7<sup>FST</sup> |
| 11 | 12 | Left ear | No/yes | 5 | - | 1 | GT | 0.5 / - | 3.4 |
| 11 | 13 | Shoulder | No/yes | 6 | > weeks | 2 | GT + FST | 0.5 / 2.5 | 7.2<sup>GT</sup> |
|  |  |  |  |  |  |  |  |  | 6.1<sup>FST</sup> |
| 12 | 14 | Left shoulder | No/yes | 4 | > weeks | 2 | GT + FST | 0.5 / 1.3 | 5.1<sup>GT</sup> |
|  |  |  |  |  |  |  |  |  | 4.8<sup>FST</sup> |
| 12 | 15 | Right shoulder | No/yes | 4.3 | > weeks | 2 | GT + FST | 0.7 / 1 | 2.7<sup>GT</sup> |
|  |  |  |  |  |  |  |  |  | 4.7<sup>FST</sup> |
| 13 | 16 | Right front leg | No/yes | 2.4x1 cm | > weeks | 2 | GT + FST | 0.5* | 3.8<sup>GT</sup> |
|  |  |  |  |  |  |  |  |  | 5.5<sup>FST</sup> |
| 14 | 17 | Hip | Yes/yes | 5x2 cm | > 3-5 days | 1 | GT | 0.1 / - | 3.9 |

### RNA extraction

All tissue samples were homogenized using Qiazol lysis buffer (Qiagen, Hilden, Germany), M-tubes (Miltenyi Biotec, Lund, Sweden) and a gentleMACS™ Octo Dissociator (Miltneyi Biotec). RNA was extracted from the homogenized tissue samples according to manufacturer’s instructions using the miRNeasy Mini Kit (Qiagen, Hilden, Germany). Additionally, all samples underwent on-column DNase digestion with the RNase-free DNase set (Zymo Research), also according to manufacturer’s instructions.

Quantity of the extracted RNA was measured using a spectrophotometer (NanoDrop ND-1000, Thermo Scientific, United States). The RNA integrity of all samples was measured on an Agilent 2100 Bioanalyzer (Agilent Technologies, Nærum, Denmark).

### cDNA synthesis

cDNA synthesis was performed from 500 ng total RNA in duplicates using the QuantiTect Reverse Transcription kit (Qiagen, Hilden, Germany) following the manufacturer’s protocol. Each of the newly synthesized cDNA samples were diluted 10 times in low EDTA TE buffer (VWR) before being stored at -20°C before pre-amplification. Additionally, non-reverse-transcriptase controls were made to assess potential DNA contamination of the samples.

### Primer design

96 pairs of primers were either chosen from an existing primer library or designed based on a panel of genes known to play a role in wound healing (See Supplementary table 1). For the new primers, the correct gene was found in Ensembl and the coding sequence was extracted. Primer3 (https://bioinfo.ut.ee/primer3-0.4.0/) was used to design primers with an amplicon size of 70-90. Whenever possible, one of the primers of the pair were designed to span an intron. When not possible, primer pairs with the two primers on different exons were chosen. All primers were synthesized by Sigma Aldrich.

### Pre-amplification

A 200nM primer mix was prepared by adding equal amounts of all primers to low-EDTA TE buffer (VWR). Pre-amplification was performed by incubating 3 µl TaqMan PreAmp Master Mix (Applied Biosystems, CA, USA), 2 µl low-EDTA TE buffer (VWR) and 2.5 µl primer mix with 2.5 µl diluted cDNA sample at 95°C for 10 minutes, then 18 cycles of 95°C for 15 seconds and 60°C for 4 minutes.

All pre-amplified samples were incubated with 4 µl of 4U/µl exonuclease (New England Biolabs) at 37°C for 30 minutes and then 80°C for 15 minutes. From all pre-amplified and exonuclease-treated samples 2 µl were taken and pooled to create a stock for the standard curves. Subsequently, all samples were diluted 10 times in low-EDTA TE buffer (VWR).

### Microfluidic qPCR

To assess the stability of the chosen 96 primer pairs, microfluidic qPCR was performed on pre-amplified cDNA made from the intentionally degraded skin and granulation tissue samples in addition to selected non-degraded experimental skin and granulation tissue samples. In total 96 samples were tested against the 96 primers in a 96.96 Dynamic Array (Fluidigm, San Francisco, CA, USA). In this array 96 samples are combined with 96 sets of primers to make 9216 individual PCR reactions in nanolitre volumes.

Of the 96 primer pairs used for the 96.96 Dynamic Array, 24 robust primers (Table 3) were chosen for qPCR on the subsequent 192.24 Dynamic Array (Fluidigm, San Francisco, CA, USA), which combines 192 samples with 24 primer pairs. The panel of 24 primers was selected based on highest percentage of valid data, i.e. fewer missing values, least amount of variation between technical cDNA replicates and primary gene function in wound healing.

**Table 3.**
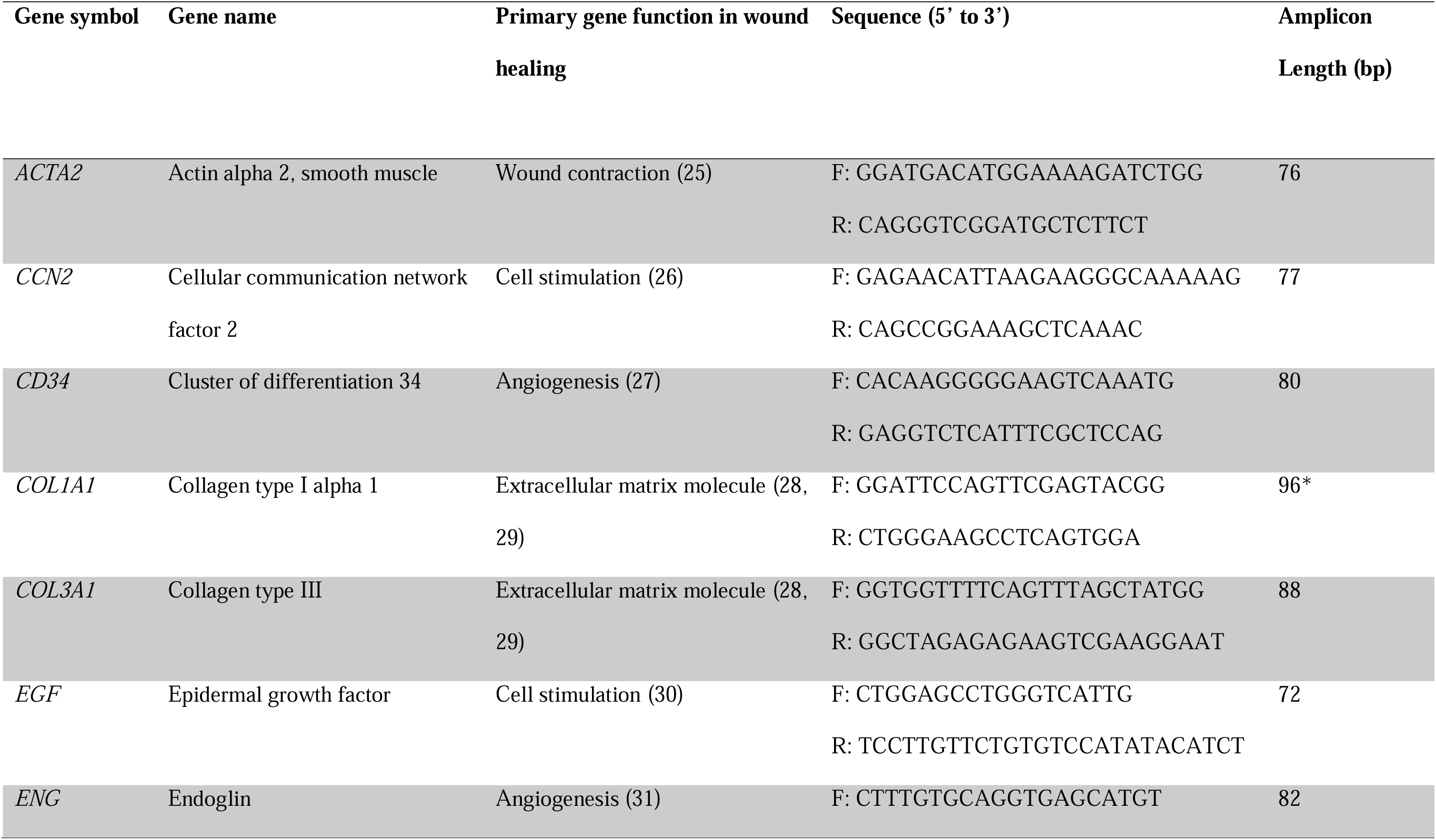
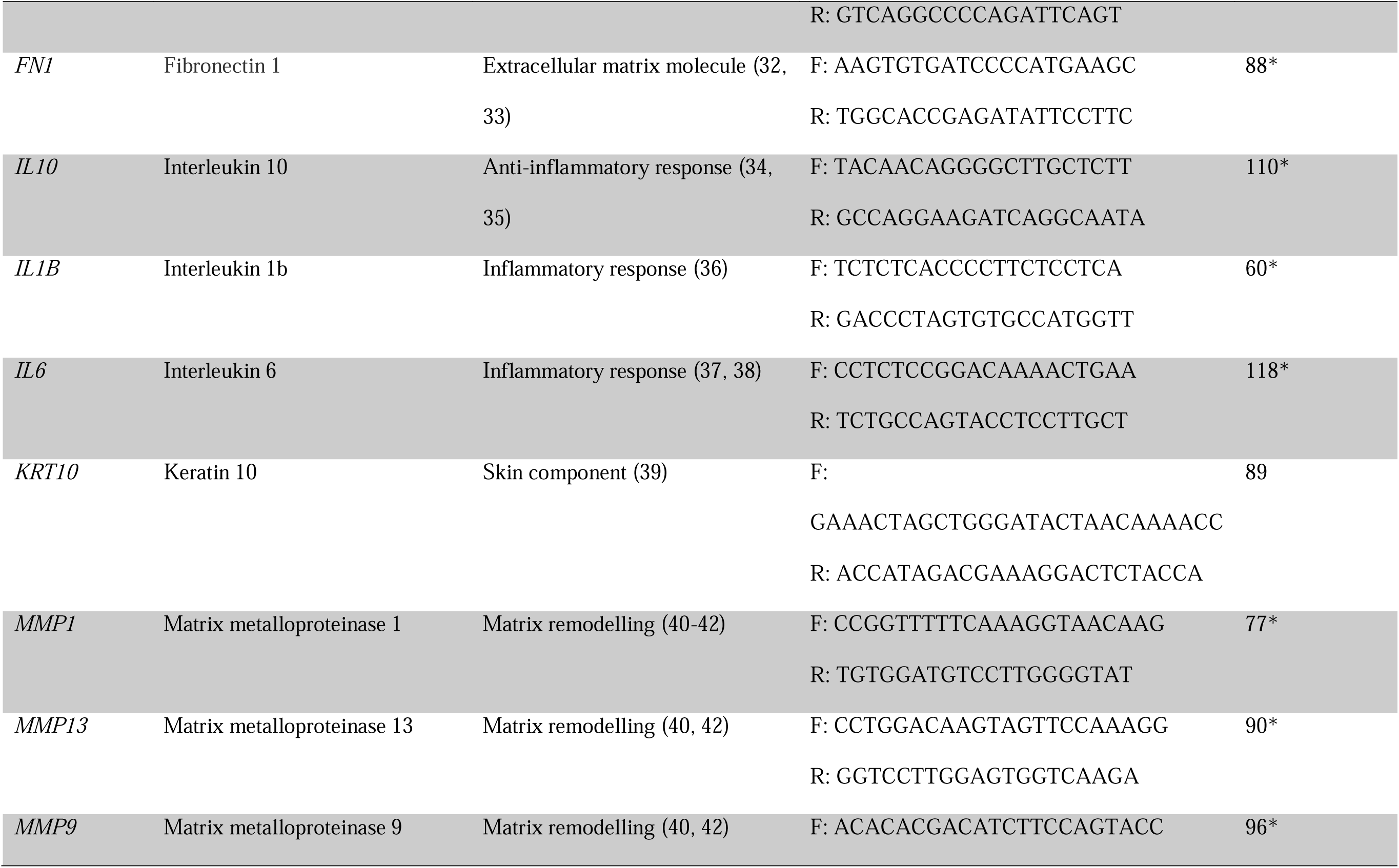
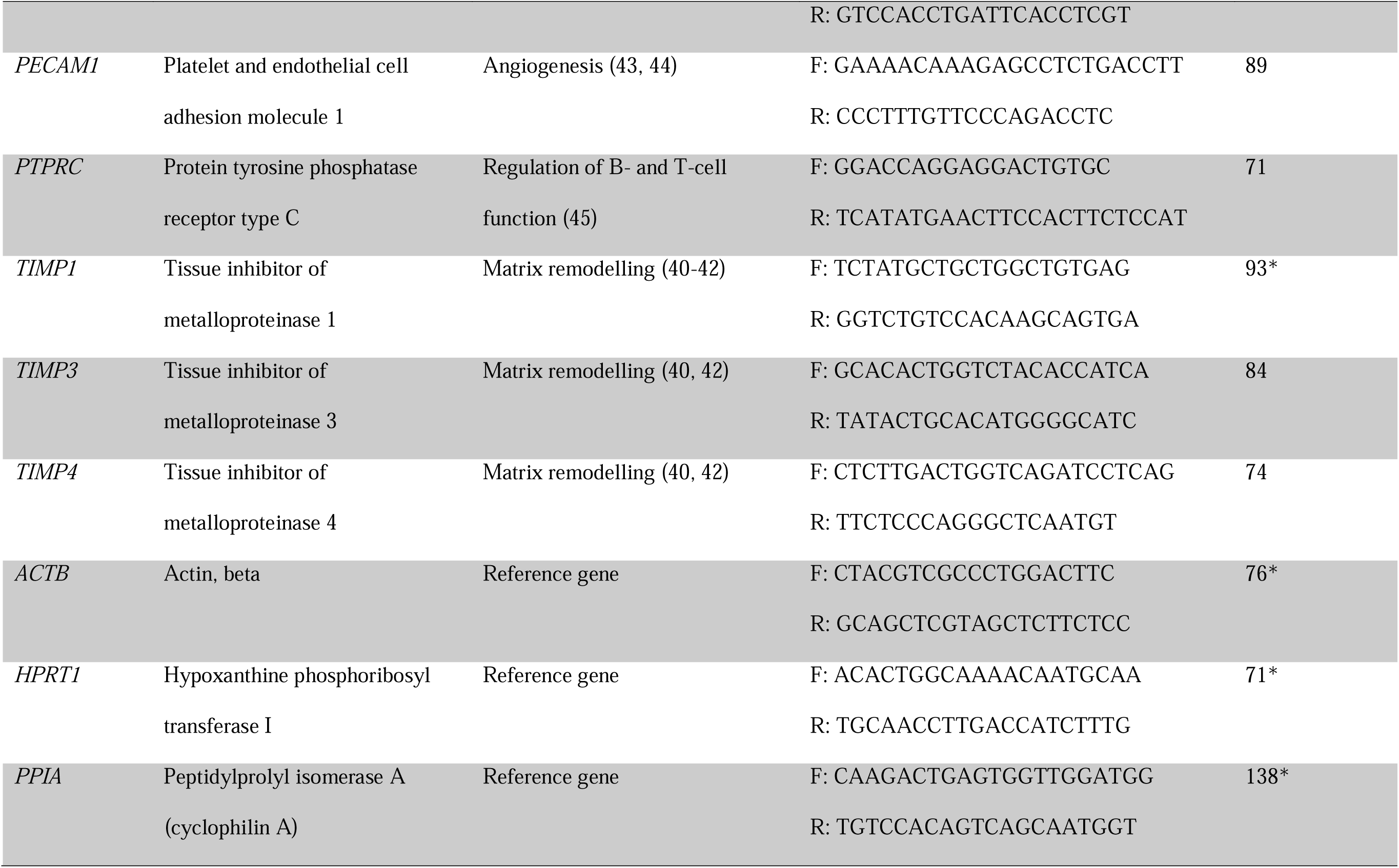
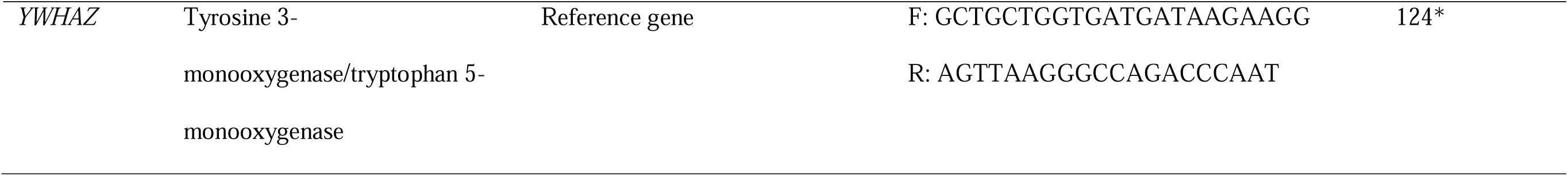
Robust primer panel of selected targets for gene expression signature. Gene symbol, gene name, gene function and sequences (5’ to 3’) of forward (F) and reverse (R) primers. 20 genes of interest were studied, and 4 reference genes were used. *Amplicon size from primer pair chosen from already existing library.

### Data analysis and statistics

To compensate for variation between the two 192.24 Dynamic Arrays, 6 of the pooled samples were used as interplate calibrators. The following described procedures were performed for both the 96.96 Dynamic Array and the 192.24 Dynamic Array. All samples were corrected for the PCR efficiency using the standard curve for each primer. Both GeNorm and NormFinder were used to identify the most stable reference genes (β-actin, HPRT1, PPIA, YWHAZ) which were then used to normalize all genes using GenEx7 (MultiD Analyses AB Sweden) in both types of arrays. Subsequently variation between technical repeats of cDNA was determined. If variation between the cDNA repeats of the experimental samples was higher than 1.5 Cq (quantification cycle) for more than 15% of a specific replicate the data for this replicate was excluded from further analysis. Due to poor RNA quality of the forensic samples, we allowed up to 20% variation for these samples. Additionally, up to 20% deviation of 1.5 Cq between replicates was allowed for each specific primer, also due to quality of the forensic samples. The average of the cDNA technical repeats was calculated before replacing the missing values (2 out of 2842, 0.07% on 192.24 Dynamic Array; 30 out of 3250, 0.92% on 96.96 Dynamic Array) with the highest Cq value measured for a certain primer added with 1. Finally relative quantities were calculated and scaled to the lowest expression (highest measured Cq value set to 1) and data was log_2_ transformed before statistical analysis.

A one-sample t-test was performed in RStudio (version 2024.12.0) to test for differences between wounds A and D within each gene. It was determined that the two wounds were not statistically significantly different, and an average of the wounds was calculated and used for further analysis.

#### Principal component analysis (PCA) was performed in RStudio

QQ plots were made in GraphPad Prism (version 10.4.1) to determine if data followed normal distribution. Data was determined to follow normal distribution for all primers. Furthermore, variance, standard deviation (SD) and ratio between the largest and smallest SD was calculated to determine which statistical test to use. An ordinary one-way ANOVA (analysis of variance) was used for primers which had equal SD, meaning the ratio was smaller than 2, and Brown-Forsythe and Welch ANOVA was used for primers which had non-equal SD. Additionally, Holm-Sidak correction and Dunnett’s T3 correction, respectively for the two ANOVA’s, were used to adjust *P* values to account for multiple comparisons.

## Results

### RNA quality

The RNA integrity number (RIN) values for the intentionally degraded RNA samples were in the range of 2.5 to 8.5, averaging at 6.7. The RIN values for the experimental samples were in the range of 6.1 to 10, averaging at 8.8. Similarly to the deliberately degraded RNA samples, the RIN values for forensic cases ranged from 2.2 to 7.2, but averaged at 4, a slightly lower mean compared to the experimentally degraded samples.

### Robust primers

Of the 96 primer pairs originally tested 24 pairs, 4 reference genes and 20 genes of interest, were chosen for further testing on the experimental and forensic samples. Due to their low percentage of missing data and reduced technical variation (Table 4) they were deemed the most stable and therefor most suitable for generating reliable results from degraded RNA in forensic samples.

**Table 4.**
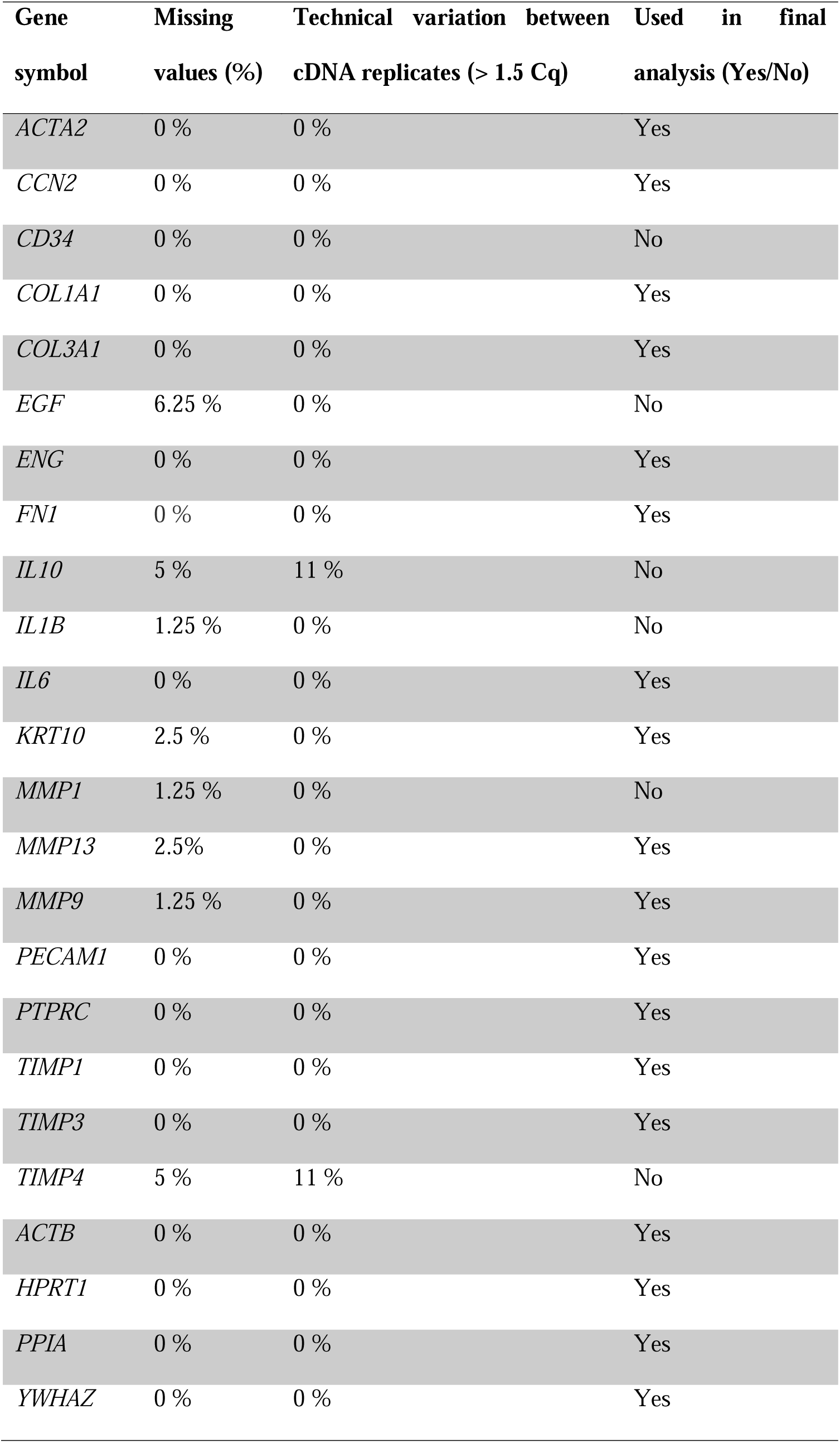
Stable primers selected based on intentionally degraded RNA. Gene symbol, percentage of missing values (out of 80) within each primer, technical variation and if the primer was used in the final analysis on all experimental and forensic samples. The technical variation indicates the percentage of technical cDNA repeats that vary from each other with more than 1.5 Cq.

### Gene expression

After manual processing of the data, 14 of the original 20 genes of interest were deemed acceptable and included in further analysis (Table 4). Due to poor RNA quality of the forensic samples and to ensure reproducibility of the analysis, all samples with a RIN score below 3 were removed from the final analysis (n = 11). The principal component analysis exhibited separate grouping of control tissue and experimental wounds of 5 days of age (Figure 2, green and yellow respectively). Additionally, wounds collected 10-15 days after wounding (orange) was seen as a cluster separated from wounds of 20-30 days of age (red). A less delimited grouping of the wounds of 35 days of age could be observed (blue). (Figure 2). The greatest divergence from the expression pattern observed in the control skin was observed in the wounds of 5 days of age. However, as the granulation tissue and wounds mature, we observe the expression pattern move towards that of the control skin.

**Fig. 2:**
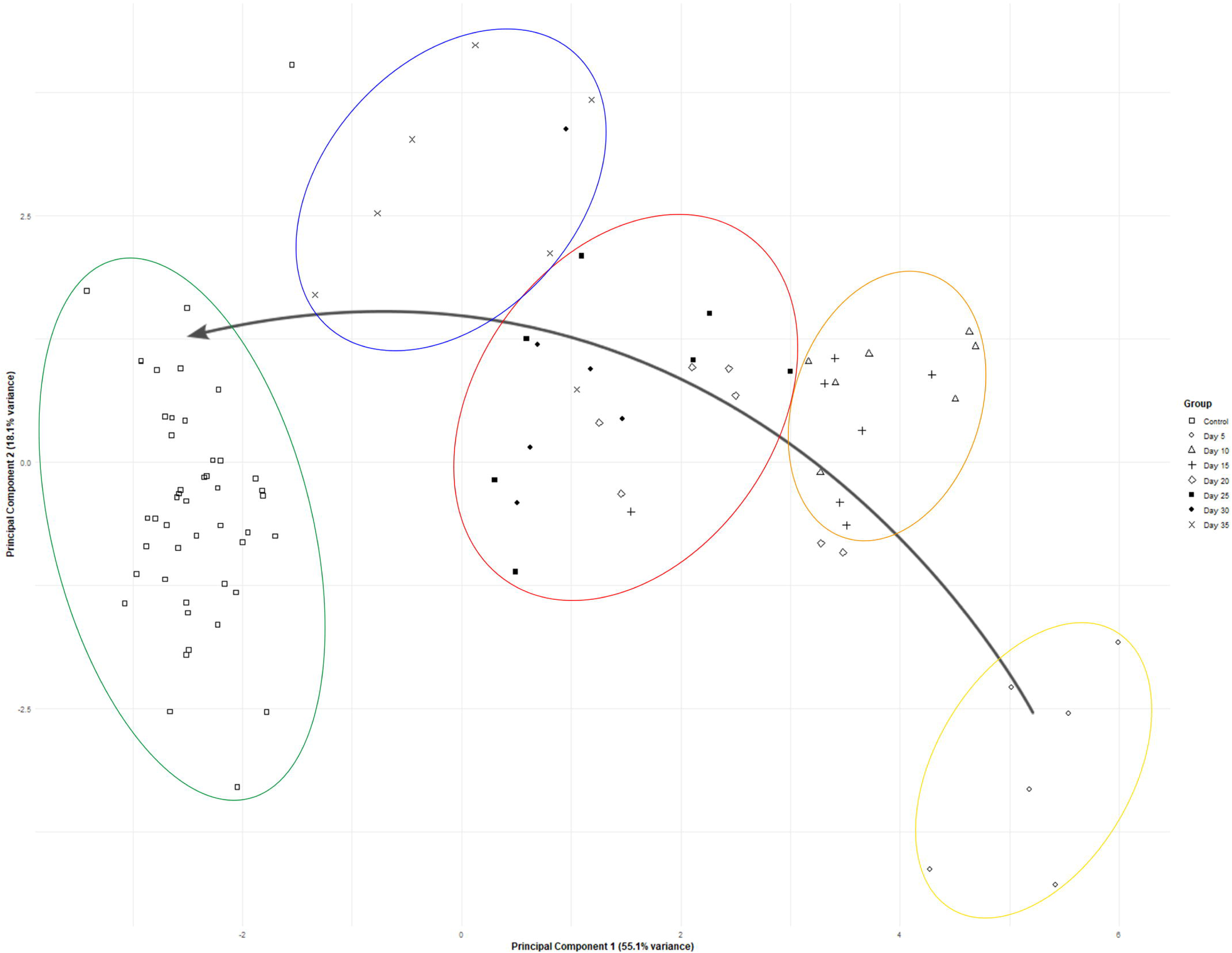
Principal component analysis (PCA) scores based on samples of control tissue and experimental granulation tissue of 5 to 35 days of age. The control samples and 5 day old wounds form two separate clusters. The 35 day old wounds form a less delimited group. The percentages in the brackets indicate the percentage of variation of data explained by the two principal components. Yellow = Day 5, orange = Day 10 and 15, red = Day 20, 25, and 30, blue = Day 35, green = Control tissue.

The loading plot extracted for the PCA of the experimental samples revealed that high expression of collagen type I (COL1A1), collagen type III (COL3A1), keratin 10 (KRT10), and cellular communication network factor 2 (CCN2, also known as connective tissue growth factor) were characteristic for the control samples and wounds of 35 day of age (Figure 3). It can also be seen that high expression of interleukin 6 (IL6), matrix metalloproteinase 9 (MMP9) and matrix metalloproteinase 13 (MMP13) were characteristic for the wounds of 5 days of age. Several of the 14 genes of interest displayed time-dependent expression (Supplementary figure 1). COL1A1, COL3A1 and KRT10 all displayed the same patterns of expression having the lowest expression at day 5 and an increase and return to the same expression levels as in the control skin and having the highest levels of expression in the older wounds (Figure 4).

**Fig. 3:**
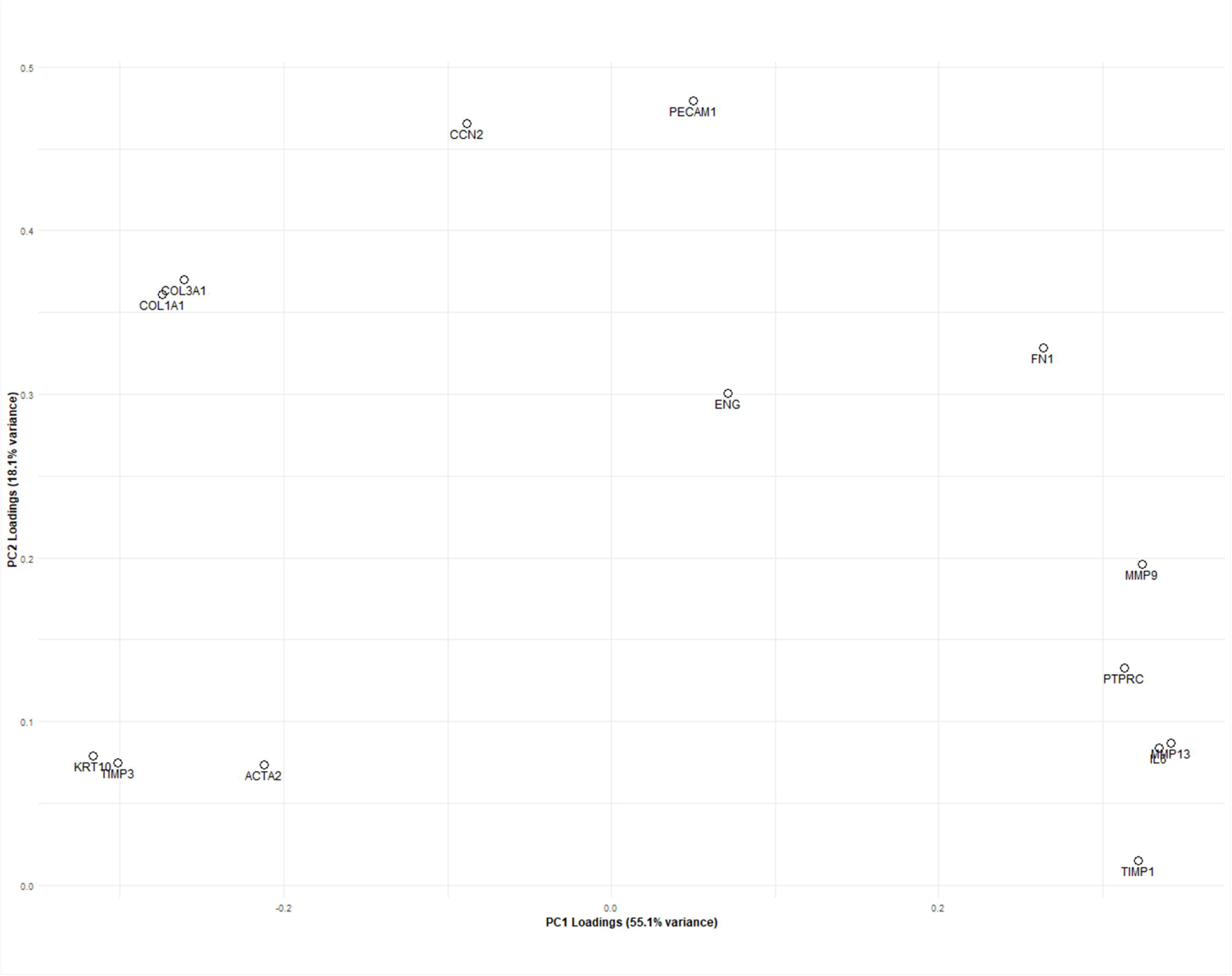
Loading plot from the principal component analysis based on the experimental samples of skin and granulation tissue. The percentages in the brackets indicate the variation of the data explained by the two principal components.

**Figure 4:**
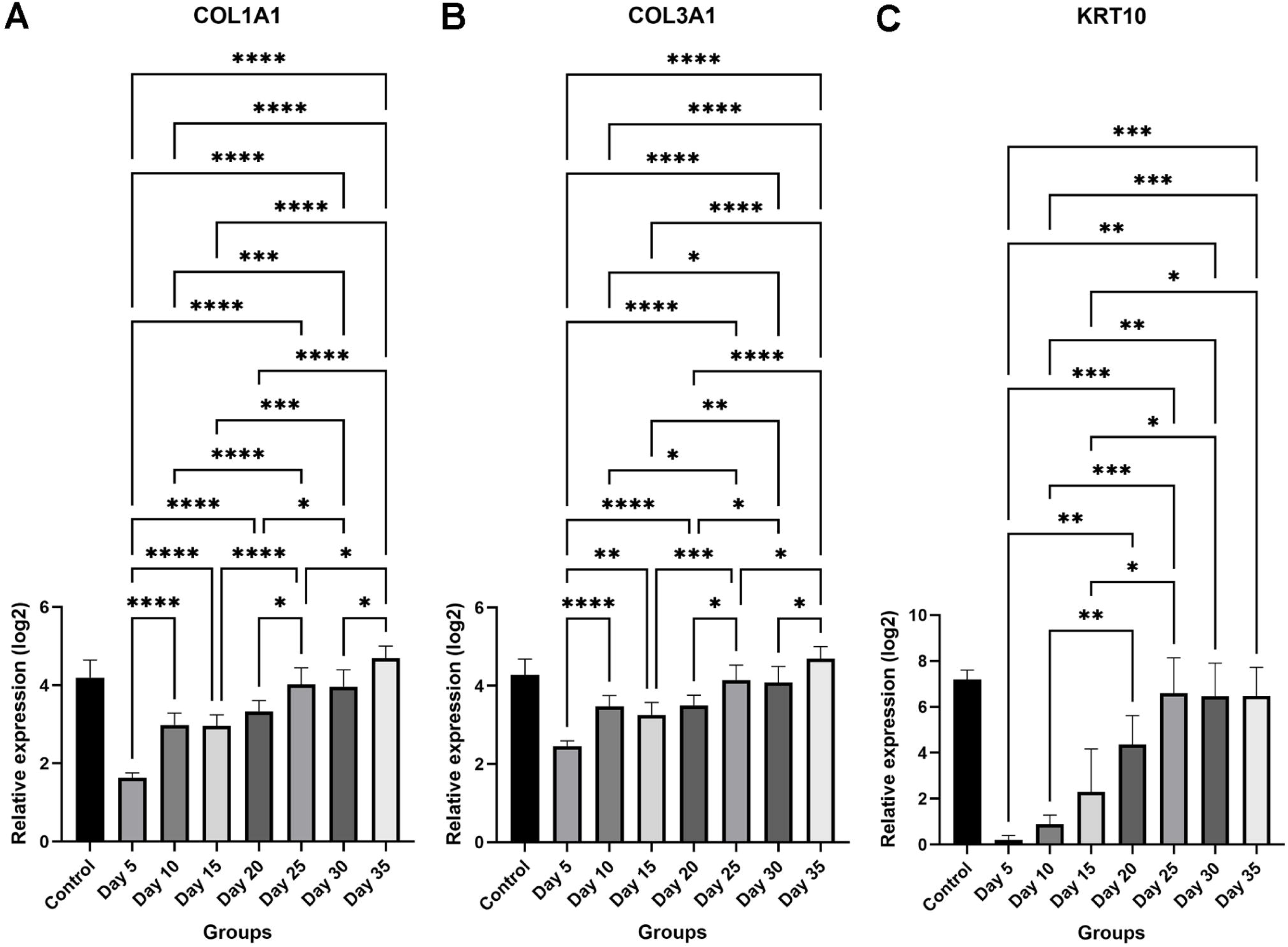
Gene expression of collagen genes and connective tissue growth factor. A) Collagen type 1 (COL1A1), B) collagen type III (COL3A1), and C) keratin 10 (KRT10). *: *P*-value = 0.01-0.05, **: *P*-value = 0.01-0.001, ***: *P*-value = 0.001-0.0001, ****: *P*-value < 0.0001.

Not surprisingly, an opposite expression pattern could be observed for the pro-inflammatory cytokine IL6, that showed a marked increase in expression which dropped with increasing wound age, however, it did not reach the same baseline level as in the control samples (Figure 5A). Similar patterns of expression were observed for the matrix metalloproteinases, which are responsible for cleaving a variety of extracellular matrix proteins, especially as a result of skin injury, but also to a limited degree in intact skin (Toriseva et al., 2012). For both MMP9 and MMP13, higher expression could be seen in younger wounds with a decrease as the wounds heal, though without reaching the levels observed in the control tissue (Figure 5B-C).

**Figure 5:**
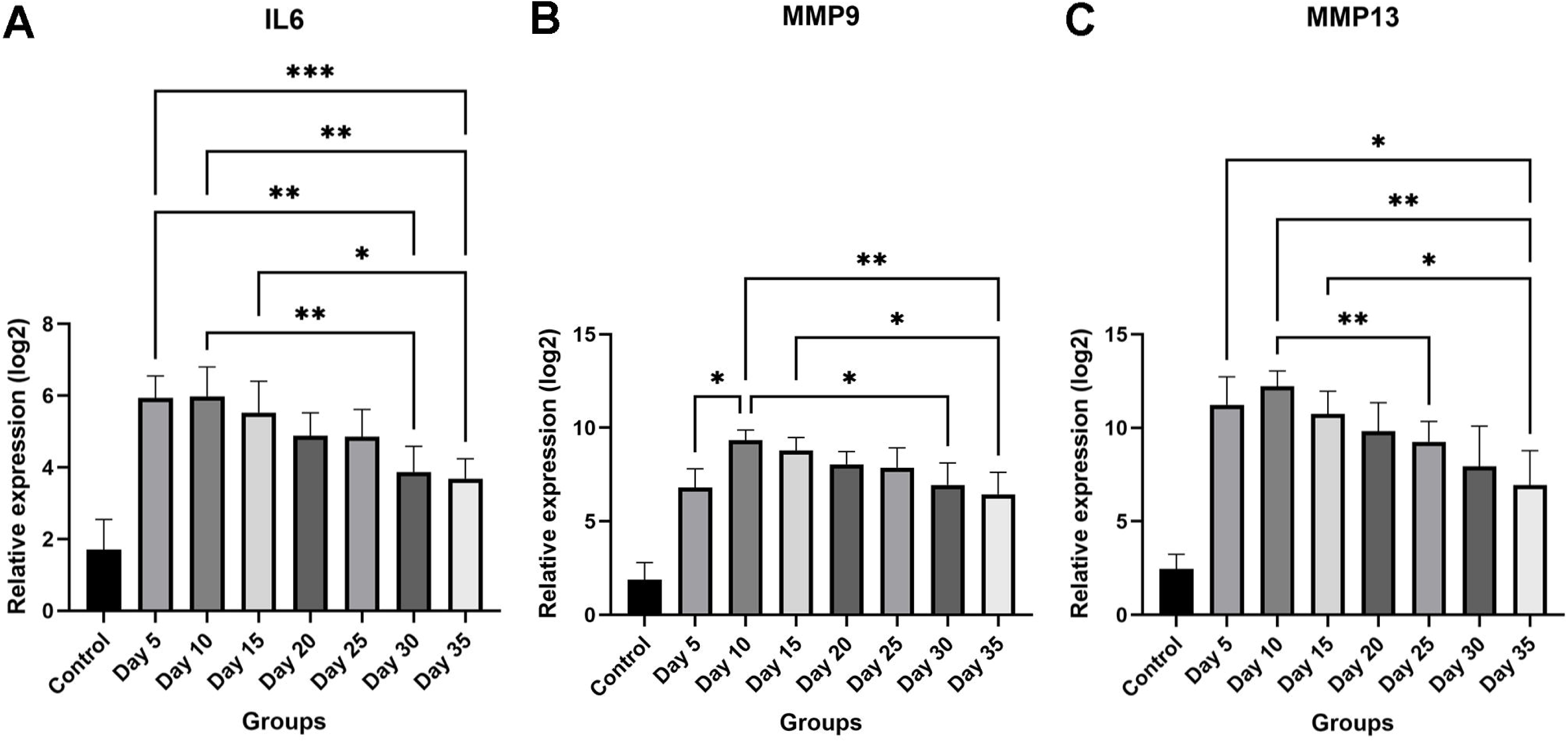
Gene expression of interleukin 6 and matrix metalloproteinases 9 and 13. A) Interleukin 6 (IL6), B) Matrix metalloproteinase 9 (MMP9), C) Matrix metalloproteinase 13 (MMP13). *: *P*-value = 0.01-0.05, **: *P*-value = 0.001-0.01, ***: *P*-value = 0.001-0.0001.

### Forensic samples

A PCA plot including experimental granulation tissue and tissue samples from forensic porcine cases is presented in Figure 6. It can be seen that some forensic samples lie close to the experimental samples while a portion of the forensic samples formed a separate cluster. The extracted loading plot displayed that the forensic samples were characterized by high expression of IL6, similarly to the younger experimental wounds, in addition to alpha smooth muscle actin (ACTA2), platelet and endothelial cell adhesion molecule 1 (PECAM1), and endoglin (ENG) (Figures 6 & 7), none of which displayed time-dependent expression patterns in the experimental samples (Supplemental figures 1C, 1D and 1F).

**Fig. 6.**
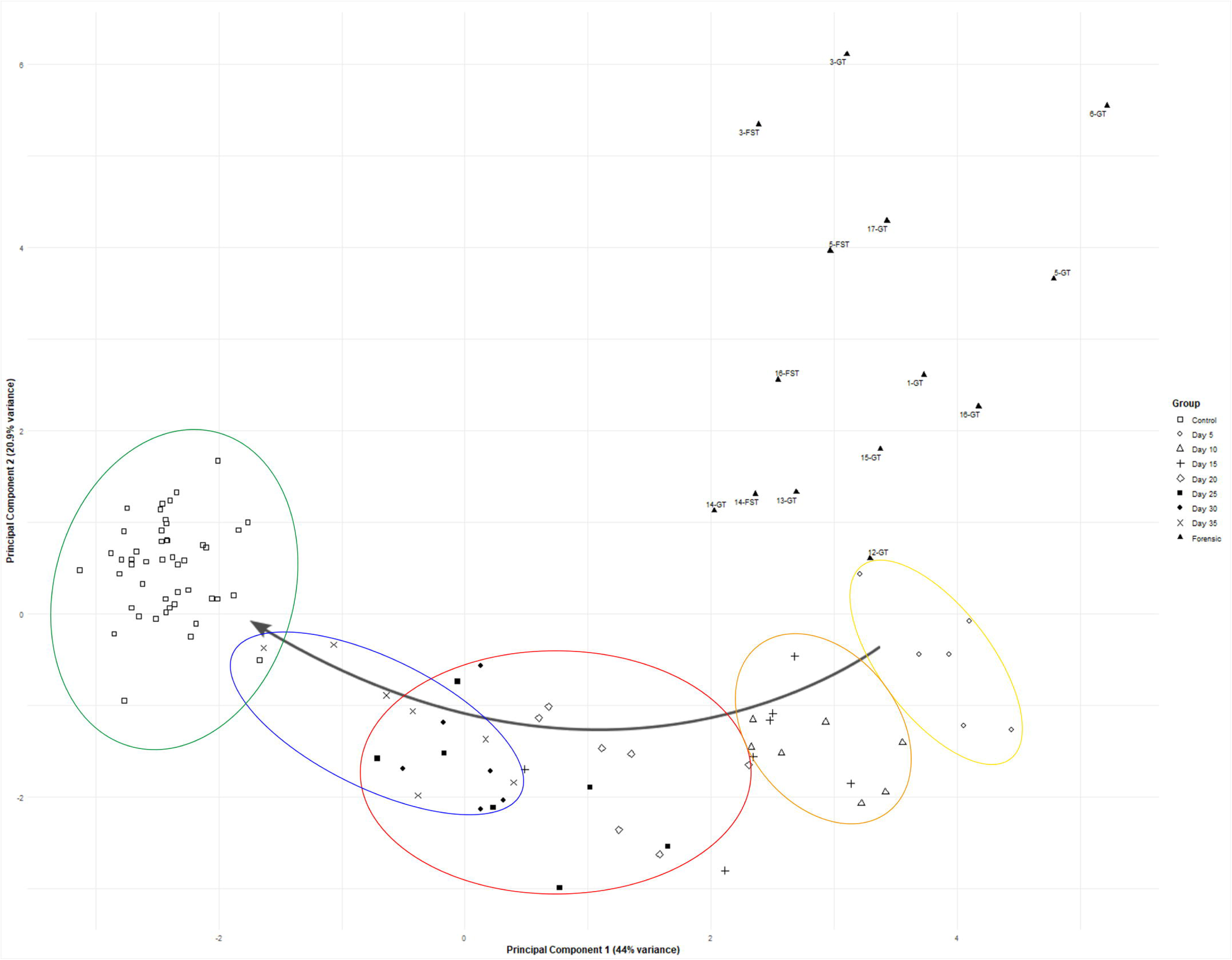
Principal component analysis (PCA) scores based on samples of control tissue, experimental granulation tissue of 5 to 35 days of age and forensic samples. Experimental samples tend to group according to age. The percentages in the brackets indicate the percentage of variation of data explained by the two principal components. Numbers of the forensic cases reference to the wound numbers listed in Table 2. GT = granulation tissue, FST = fibrous scar tissue. Yellow = Day 5, orange = Day 10 and 15, red = Day 20, 25, and 30, blue = Day 35, green = Control tissue, Black triangles = Forensic samples.

**Fig. 7:**
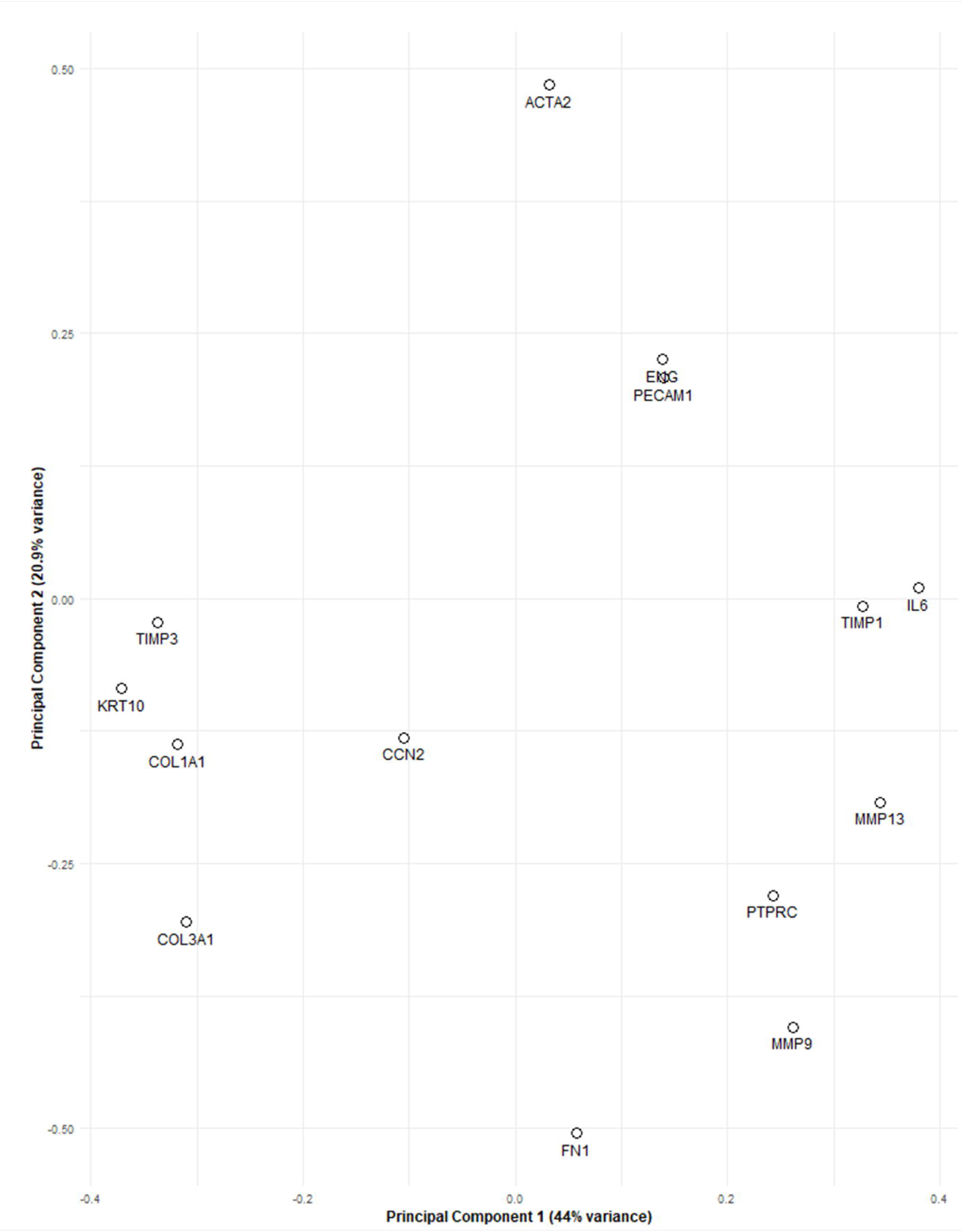
Loading plot from the principal component analysis based on the experimental samples of skin and granulation tissue and forensic samples. The percentages in the brackets indicate the variation of the data explained by the two principal components.

## Discussion

The present study demonstrated a time-dependent gene expression profile in experimental porcine granulation tissue. The PCA based on data from 14 highly robust primers, revealed that the experimental samples displayed a pattern of grouping according to wound age. At day 5, the samples display the greatest divergence from the controls, however, as the wound age increases the granulation tissue matures, and the gene expression gradually returns towards the levels expressed in intact skin.

The primer pairs included in the study were selected based on their roles in the wound healing process. The wound healing process is comprised of several overlapping phases: haemostasis, inflammation, proliferation and remodelling (Huang et al., 2022, Kondo, 2007, Shah et al., 2012, Wang et al., 2018). IL6 is found early in wound healing and is known to elicit both pro-and anti-inflammatory effects (Johnson et al., 2020). Indeed, marked increase of IL6 expression was observed in the youngest wounds with a continuous drop in expression levels as wound age increased.

The proliferative phase, which start at 2-3 days after wounding and can last for 2-3 weeks is characterized by fibroblast infiltration, production of immature collagen, primarily type III, and angiogenesis (Almadani et al., 2021). In the present study mRNA expression levels of collagens I and III were found at the lowest levels of expression in the wounds of 5 days of age with a steady increase in expression with increasing wound age. In a murine excisional wound model, time dependent levels of collagens I and III were likewise observed, with collagen III being the dominant type early in the wound healing process and collagen I being dominant later in the wound healing process (Wang et al., 2016). Collagens, however, are known to undergo posttranslational modifications, such as phosphorylation, acetylation, and ubiquitination (Guo et al., 2025, Kuivaniemi and Tromp, 2019), which could explain the discrepancy between our mRNA expression patterns and the protein levels observed by Wang et al (Wang et al., 2016). Similar to the present study, an excisional porcine wound model observed low expression levels for both collagen I and III early in wound healing with an increase during the later phases (Ulrich et al., 2007).

The last phase, the remodelling phase, is characterised by the substitution of granulation tissue with fibrous scar tissue as collagen III is replaced by collagen I (Almadani et al., 2021). During this phase and the previous phases, a continuous degradation and production of collagen occurs. The temporary, immature collagen is broken down and substituted with permanent fibrous scar tissue rich in collagen type I (Singh et al., 2023). The MMPs are responsible for the breakdown of collagens, and other extracellular matrix proteins, while the tissue inhibitor of metalloproteinases (TIMPs) inhibit the effects of MMPs to ensure a balanced level of collagen breakdown and deposition (Murphy and Nagase, 2008, Nagase et al., 2006). MMP9 and MMP13 were both expressed in higher levels in the younger wounds, with a slight decrease in expression as wound age increased. However, expression levels did not reach the baseline level of the control skin, indicating a continuous degradation of collagen in granulation tissue at day 35. In a porcine wound model with sterile dressings, a similar expression pattern of MMP9 was observed (Wang et al., 2000). As would be expected during a balanced wound healing, TIMP1 displayed a similar pattern of expression with higher levels in younger wounds and a gradual decrease throughout the wound healing process (Supplementary figure 1A). This pattern of MMP9 and TIMP1 expression with highest levels in the younger wounds and a subsequent decrease of expression has also been observed in previous studies in pigs and rats (Kananykhina et al., 2020, Ulrich et al., 2007).

Myofibroblasts, which express alpha-smooth muscle actin (ACTA2) play an important role in contraction of a wound (Desmouliere et al., 2005). However, in murine experimental wounds myofibroblasts contributed to wound contraction but were not necessary for successful wound healing (Ibrahim et al., 2015). In an immunohistochemical study of experimental porcine wounds, the numbers of alpha-smooth muscle actin positive myofibroblast decreased with increasing wound age (4). Therefore, we would have expected to observe a time-dependent expression of ACTA2 in the granulation tissue, however, this was not the case (Supplementary figure 1C). The discrepancy between the mRNA expression pattern we observed, and protein expression pattern observed by Pankoke et al (Pankoke et al., 2023) could potentially be explained by post translational modifications. Even if there is a continuous production of ACTA2 mRNA, protein levels could still reflect a time-dependent pattern.

Besides being chosen based on their role in wound healing, all primers were tested for their stability on intentionally degraded RNA. Although all 20 selected primer pairs were tested on intentionally degraded samples beforehand, only 14 met the quality control criteria and were deemed suitable for statistical analysis of the forensic samples. The RNA quality of the forensic samples proved even poorer than that of the intentionally degraded samples. This could be due to the way the forensic cases were handled prior to sampling. The tissue, whole carcasses or body parts, were frozen and subsequently thawed at room temperature prior to forensic evaluation. Furthermore, in 9 out of 14 cases the carcases were scalded, singed and scraped as a part of the slaughter process, before tissue sampling (Barington and Jensen, 2013). This process removes the outermost layers of the skin, potentially affecting RNA quality and the comparability to our experimental samples.

The addition of the forensic samples to the PCA (Figure 6) shows that the forensic cases cluster on their own. However, we do see an overlap between the experimental samples of ages 5-15 days and the forensic samples on principle component 1. There appear to be no specific sub-grouping of the forensic cases that have undergone slaughter treatment from cases that have not. This difference between the forensic samples and the experimental could be the result of many different influences. The poor RNA quality of the forensic samples as a result of the current sampling procedures could potentially affect their comparability to the experimental samples. Additionally, the pigs in the forensic cases were mainly slaughter pigs weighing approximately 90 kg and sows that typically weigh up to 200-300 kg. Therefore, the pigs from the forensic cases were older, as well as larger than the experimental pigs included in this study. Previous studies in murine models have observed delayed cutaneous wound healing in older individuals compared to younger individuals (Jeong et al., 2008, Plum et al., 2025). However, as slaughter pigs are only 5-6 months old when they reach slaughter it seems unlikely that the age of the pigs is the main reason for the difference between the forensic and experimental wounds. The sex of the slaughter pigs were unknown. The biological variation, sex of the pigs, wound size and location of the wound on the body could contribute to the difference between experimental samples and part of the forensic samples. Finally, it possible that the difference in environment between the industrial pigs included as forensic cases and the experimental pigs might also add to the difference observed in their wound healing patterns.

The PCA plot of the experimental and forensic samples and its corresponding loading plot also showed that many of the forensic samples appeared to be characterized by higher expression of IL6, similarly to the younger wounds. In addition, the forensic samples were also characterized by higher expression of ACTA2, which plays a role in wound contraction (Ibrahim et al., 2015), ENG, and PECAM1, both of which play a role in angiogenesis (Valluru et al., 2011, Vasalou et al., 2023, Watt et al., 1995). None of these three displayed time-dependent expression in the experimental wounds. This pattern of higher expression of ACTA2, ENG, and PECAM1, combined with the increased thickness of granulation tissue, compared to the experimental wounds, indicates the forensic wounds could be older than 5-15 days, despite overlapping with the experimental samples of that age on principal component 1. If the forensic wounds do indeed exceed 15 days of age, the high expression of IL6, an inflammatory marker usually found early in wound healing, could potentially be explained by repeated trauma. It is also possible that the forensic samples do not group within experimental samples because their age exceeds 35 days. Only a few of the forensic wounds were estimated to be younger than one week. The rest were estimated to be several weeks of age. No grouping based on the estimated wound age was observed among the forensic samples. This may be due to the age estimated being too imprecise. However, the true age of the wounds is unknown. Moreover, a primer panel of only 14 primers may not be large enough to age determine forensic samples. A larger primer panel could potentially detect gene expression patterns that the present primer panel did not.

## Conclusion

It can be determined from this study that it is possible to design primers that can be used on highly degraded RNA and that these primers can be used to establish a time-dependent gene expression profile in a porcine wound model. However, the current procedure of storing and handling of forensic cases before sampling has a negative impact on RNA quality. In order to implement gene profiles for age assessment of wounds in forensic cases new sampling protocols must be implemented. Finally, qPCR as a sole method is not sufficient for precise age estimations. Additional methods of investigation must be used concurrently.

## Statement of author contributions

**C. Bækgård**: Conceptualization, Data curation, Formal analysis, Investigation, Methodology, Project administration, Visualization, Writing – original draft. **K. Skovgaard**: Methodology, Investigation, Supervision, Writing – review & editing. **M.H. Hansen**: Writing – review & editing. **H. E. Jensen**: Conceptualization, Writing – review & editing. **K. Barington**: Conceptualization, Methodology, Funding acquisition, Project administration, Supervision, Investigation, Writing – review & editing.

## Acknowledgements

The authors would like to thank Karin Tarp for assistance in lab work. We also acknowledge Dennis Brok and Frederik Andersen for technical assistance. Finally, we would like to thank Karen Pankoke for collecting granulation tissue from the forensic cases.

## Ethical statement

The study and procedures were approved by the Danish Animal Inspectorate

---

**Supplementary fig. 1** Gene expression of A) Tissue inhibitor of metalloproteinase 1 (TIMP1), B) Tissue inhibitor of metalloproteinase 3 (TIMP3), C) Alpha-smooth muscle actin (ACTA2), D) Platelet and endothelial cell adhesion molecule 1 (PECAM1, also known as CD31), E) Protein tyrosine phosphatase receptor type C (PTPRC, also known as CD45), F) Endoglin (ENG, also known as CD105), G), Cellular communication network factor 2 (CCN2, also known as Connective tissue growth factor (CTGF)) H) Fibronectin (FN1). *: *P*-value = 0.01-0.05, **: *P*-value = 0.01-0.001, ***: *P*-value = 0.001-0.0001, ****: *P*-value < 0.0001.

