## Supplementary Table 1 for "Gene expression profiling for forensic age assessment of porcine skin wounds"

**Supplementary table 1: Primer panel**

| **Gene symbol** | **Gene name** | **Primary gene function** | **Sequence (5’ to 3’)** |
| --- | --- | --- | --- |
| ACTA2 | Actin Alpha 2, Smooth Muscle | Re-epithelialization | F: GACGTACGACTGGCATTGTG  R: GCGTAGCCCTCGTAAATGG |
| ACTA2 | Actin Alpha 2, Smooth Muscle | Re-epithelialization | F: GGATGACATGGAAAAGATCTGG  R: CAGGGTCGGATGCTCTTCT |
| ANGPT2 | Angiopoietin-2 | Angiogenesis | F: GAGGAGACCAAGGCTTACTGC  R: TCTTCACGTCGCTGAATAACTG |
| BGN | Biglycan | Extracellular matrix molecule | F: TCTCTGAGGCCAAGCTCACT  R: GTGGTCCAGATGGAGTTCGT |
| CCN2 | Cellular Communication Network Factor 2 | Cell stimulation | F: GAGAACATTAAGAAGGGCAAAAAG  R: CAGCCGGAAAGCTCAAAC |
| CCN2 | Cellular Communication Network Factor 2 | Cell stimulation | F: AGTGTGCACAGCCAAAGATG  R: GCAGCTGCTCTGGAAGGA |
| CD34 | Cluster of differentiation 34 | Angiogenesis | F: CCAACGGAACAGAACTTAGCA  R: TTCGAGAAGTTTTGGATGCTC |
| CD34 | Cluster of differentiation 34 | Angiogenesis | F: CACAAGGGGGAAGTCAAATG  R: GAGGTCTCATTTCGCTCCAG |
| COL1A1 | Collagen type I alpha 1 | Extracellular matrix molecule | F: AACATGGAGACAGGCGAGAC  R: TCTTGTCCTTGGGGTTCTTG |
| COL1A1 | Collagen type I alpha 1 | Extracellular matrix molecule | F: GGATTCCAGTTCGAGTACGG  R: CTGGGAAGCCTCAGTGGA |
| COL3A1 | Collagen type III | Extracellular matrix molecule | F: GGATGGTTGCACTAAACACACT  R: CTACAATAGGTAGTCTCACAGCCTTG |
| COL3A1 | Collagen type III | Extracellular matrix molecule | F: GGTGGTTTTCAGTTTAGCTATGG  R: GGCTAGAGAGAAGTCGAAGGAAT |
| COL4A1 | Collagen type IV alpha 1 | Extracellular matrix molecule | F: GGTGAAAAAGGCGACCAC  R: GACCCACATCTCCCTTATCG |
| COL4A1 | Collagen type IV alpha 1 | Extracellular matrix molecule | F: CCCAAAGGTGTTGATGGTTT  R: ACCTGGGTTGCCAGGTAAG |
| COL5A1 | Collagen type V alpha 1 | Extracellular matrix molecule | F: ACTTCCCGGACGGTGAATA  R: AAGTTGCAGTAGACTTTGAAGGAGTC |
| COL5A1 | Collagen type V alpha 1 | Extracellular matrix molecule | F: GTGGGAGAGAAGGGTGAGC  R: TCTCCTCTTTCGCCTTTGG |
| COL6A1 | Collagen type VI alpha 1 | Extracellular matrix molecule | F: CTGCCCGATCACCTTCTC  R: TTGGTGGTGTCGAAGTTGTG |
| COL6A1 | Collagen type VI alpha 1 | Extracellular matrix molecule | F: GATCTGCATAGACAAGAAGTGTCC  R: TCCAGCAGGATGGTGATGT |
| COL7A1 | Collagen type VII alpha 1 | Extracellular matrix molecule | F: GAGGAGGAAGGCCAGGTG  R: GGGAGCCTCAGGGTTCTG |
| COL7A1 | Collagen type VII alpha 1 | Extracellular matrix molecule | F: AAGGGAGATAAGGGAGAAGCTG  R: ATACCTGGCTCCCCCAACT |
| CXCL8 | Interleukin 8 | Neutrophil chemotaxis | F: GTACCGGACCTTCCACAAGA  R: AAGTAGCTCCCCGACACTCA |
| EGF | Epidermal growth factor | Cell stimulation | F: CTGGAGCCTGGGTCATTG  R: TCCTTGTTCTGTGTCCATATACATCT |
| EGF | Epidermal growth factor | Cell stimulation | F: CTATGCCTGCAAGACTCAGAAG  R: CAGGCCTGATACCACTCACA |
| ENG | Endoglin | Angiogenesis | F: CTTTGTGCAGGTGAGCATGT  R: GTCAGGCCCCAGATTCAGT |
| ENG | Endoglin | Angiogenesis | F: CAGTAACATCCAAGGCGTCA  R: ATGCTATCACACTGGCGTTG |
| FGF1 | Fibroblast growth factor 1 | Cell regulation | F: TTGTACGGCTCACAGACACC  R: GCGTGCTTCTTGGATGTGTA |
| FGF1 | Fibroblast growth factor 1 | Cell regulation | F: TTCCTCAGGATCCTTCCAGAT  R: GCAGCTGAATGTGCTGGTC |
| FGF4 | Fibroblast growth factor 4 | Proliferation of epidermal stem cells | F: CTCTATGGCTCGGCTTTCTTC  R: GCACTCGTAGGCGTTGTAGTT |
| FGF4 | Fibroblast growth factor 4 | Proliferation of epidermal stem cells | F: CACGAGTGACAGCCTACTGG  R: CTCATGGCCACGAAGAATC |
| FGF7 | Fibroblast growth factor 7 | Stimulation of keratinocytes | F: GGAAAACTCTATGCAAAGAAAGAATG  R: ATACGTGTTGTAATGGTTTTCCAGA |
| FGF7 | Fibroblast growth factor 7 | Stimulation of keratinocytes | F: AAGAACAATTACAACATCATGGAAATC  R: GCAAGATAATATTCACTTTCCACTCC |
| FMOD | Fibromodulin | Extracellular matrix molecule | F: CAACACCAACCTGGAGAACC  R: CACGGTGCAGAAACTGCTAA |
| FMOD | Fibromodulin | Extracellular matrix molecule | F: TCCCACAACAGTCTCACCAA  R: GAGAGGTCGAGCTCAAGGAG |
| FN1 | Fibronectin 1 | Extracellular matrix molecule | F: AAGTGTGATCCCCATGAAGC  R: TGGCACCGAGATATTCCTTC |
| HAS1 | Hyaluronan synthase 1 | Hyaluronan production | F: CAGTGGTCCCCTAGGTCTGTA  R: GCCCAGGAACTTCTGATTGT |
| HAS1 | Hyaluronan synthase 1 | Hyaluronan production | F: ACTACGTGCAGGTCTGTGACTC  R: GGGGGTCCTCGTCCAGTA |
| HAS2 | Hyaluronan synthase 2 | Hyaluronan production | F: CTCCTGGGTGGTGTGATTTT  R: TGCATAGAGCAACGTTCCAA |
| HGF | Hepatocyte growth factor | Proliferation of epidermal stem cells | F: ACCATGCGAGGGAGATTATG  R: ACCAGGAACAATGACACCAAGTA |
| HGF | Hepatocyte growth factor | Proliferation of epidermal stem cells | F: GTTTCCCTTCTCGAAACAAAGAC  R: TCTCCTCTTCCATGGACATCA |
| HIF1A | Hypoxia-induced factor-1α | Aids vascular cell proliferation | F: CCCTCTGACTTAGCCTGTAGACTC  R: GCATTAACTTCACAATCATAACTGG |
| HIF1A | Hypoxia-induced factor-1α | Aids vascular cell proliferation | F: AAGCAGTAGGAATTGGAACATTATTAC  R: TGCATCCTTTTACACGTTTCC |
| IFNA1 | Interferon alpha 1 | Activation of macrophages, inhibition of fibroblast proliferation | F: CATGAGATGCTCCAGCAGAC  R: AGCTGCTGATCCAGTCCAGT |
| IFNB1 | Interferon beta 1 | Activation of macrophages, inhibition of fibroblast proliferation | F: AACCGTCATTAAGACTATCCTTGTG  R: CTCCTCCATGATTTCCTCCA |
| IFNG | Interferon gamma | Activation of macrophages, inhibition of fibroblast proliferation | F: CCATTCAAAGGAGCATGGAT  R: TTCAGTTTCCCAGAGCTACCA |
| IL10 | Interleukin 10 | Anti-inflammatory response | F: TACAACAGGGGCTTGCTCTT  R: GCCAGGAAGATCAGGCAATA |
| IL1A | Interleukin 1a | Inflammatory response | F: TGTGCTAAATAACCTGGATGAGG  R: GGTTCGTCTTCGTTTTGAGC |
| IL1B | Interleukin 1b | Inflammatory response | F: TCTCTCACCCCTTCTCCTCA  R: GACCCTAGTGTGCCATGGTT |
| IL2 | Interleukin 2 | Inflammatory response | F: AAAGCTCTGGAGGGAGTGC  R: TGTTTCAGATCCCTTTAGTTCCA |
| IL6 | Interleukin 6 | Inflammatory response | F: CCTCTCCGGACAAAACTGAA  R: TCTGCCAGTACCTCCTTGCT |
| KRT1 | Keratin 1 | Skin component | F: CGTGAGTGTGTCTGTGAGCA  R: CAGAGCCACCACCTCCTC |
| KRT1 | Keratin 1 | Skin component | F: GAGAAGAAAGCAGGATGTCTGG  R: GATACTGGTATGGCTGGTGCT |
| KRT10 | Keratin 10 | Skin component | F: GAAACTAGCTGGGATACTAACAAAACC  R: ACCATAGACGAAAGGACTCTACCA |
| KRT10 | Keratin 10 | Skin component | F: GTTCAATGAAAAGAGCAAGGAACT  R: TCAGATTTATAGCTGGACACCTGTT |
| KRT5 | Keratin 5 | Keratinocyte attachment | F: CGAGGAGTGCAGATTGAGTG  R: AGGAGAGGGTGTTTGTGACG |
| KRT5 | Keratin 5 | Keratinocyte attachment | F: GTACCAGACCAAGTACGAAGAGC  R: GCTTGGTGTTGCGGAGAT |
| MMP1 | Matrix metallopeptidase 1 | Matrix remodelling | F: ACGAATGCTGGAGGTATGATG  R: GTTGCCAATCCCAGGAAAT |
| MMP1 | Matrix metallopeptidase 1 | Matrix remodelling | F: CCGGTTTTTCAAAGGTAACAAG  R: TGTGGATGTCCTTGGGGTAT |
| MMP13 | Matrix metallopeptidase 13 | Matrix remodelling | F: CCTGGACAAGTAGTTCCAAAGG  R: GGTCCTTGGAGTGGTCAAGA |
| MMP2 | Matrix metallopeptidase 2 | Matrix remodelling | F: TCGCTGGAGATAAGTTCTGGAG  R: GGCGTCTGCAATGAGCTT |
| MMP3 | Matrix metallopeptidase 3 | Matrix remodelling | F: GAAGCATTTGGGTTTTTCTATTTC  R: ACTTTCTTTGCATTTGGGTCA |
| MMP3 | Matrix metallopeptidase 3 | Matrix remodelling | F: GGACAAATACTGGAGATTTGATGAG  R: GGTTCAACCCCTGGAAAGTC |
| MMP8 | Matrix metallopeptidase 8 | Matrix remodelling | F: GGCTGCCTATGAGGATTCTG  R: TGAATGTCATAGCCGCTCAG |
| MMP9 | Matrix metallopeptidase 9 | Matrix remodelling | F: ACACACGACATCTTCCAGTACC  R: GTCCACCTGATTCACCTCGT |
| PDGFA | Platelet-derived growth factor subunit A | Cell chemotaxis | F: AACACCAGCAGCGTCAAGT  R: TTCCTGACGTATTCCACCTTG |
| PDGFB | Platelet-derived growth factor subunit B | Cell chemotaxis | F: CGTCCAGGTGAGAAAGATCG  R: AGGTGGTCCTCCAAGGTCAC |
| PECAM1 | Platelet and endothelial cell adhesion molecule 1 | Angiogenesis | F: GAAAACAAAGAGCCTCTGACCTT  R: CCCTTTGTTCCCAGACCTC |
| PECAM1 | Platelet and endothelial cell adhesion molecule 1 | Angiogenesis | F: CAAGGTGATAGCCCCAGTG  R: CTGCCCAGACTCCACCTC |
| PTPRC | Protein Tyrosine Phosphatase Receptor Type C | Regulation of B- and T-cell function | F: CCTACACTTGAGCAGTACCAATTC  R: TTCTTTACTTGTCCGTTCTGAGC |
| PTPRC | Protein Tyrosine Phosphatase Receptor Type C | Regulation of B- and T-cell function | F: GGACCAGGAGGACTGTGC  R: TCATATGAACTTCCACTTCTCCAT |
| TBXAS1 | Thromboxane A synthase 1 | Vasoconstriction | F: CTCGTTTTAATCCTATCATTTCCATC  R: CTCGCTTCTTATTGGGCAAA |
| TBXAS1 | Thromboxane A synthase 1 | Vasoconstriction | F: GACATCCAGAGGTGCTACTGCT  R: GCTCCTCGCTGGAGTTCA |
| TGFA | Transforming growth factor-α | Mitogenic for keratinocytes and fibroblasts | F: TGGTGGTGGTCTCCATAGTG  R: TCACAGTGTTTTCGGACCTG |
| TGFA | Transforming growth factor-α | Mitogenic for keratinocytes and fibroblasts | F: CCCAGATTCCCACAGTCAGT  R: CAGAGTGGCAGACACATGCT |
| TGFB1 | Transforming growth factor-β1 | Cell chemotaxis and angiogenesis | F: TCACCGGGGCTGTATTTAAG  R: AAGGAAGACCCCAGTCAGGT |
| TGFB2 | Transforming growth factor-β2 | Cell chemotaxis and angiogenesis | F: GACCCCACATCTCCTGCTAA  R: ATAGGCTGCATCCAAAGCAC |
| TGFB3 | Transforming growth factor-β3 | Cell chemotaxis and angiogenesis | F: ACTGCTTCCGCAATTTGG  R: CACTTCCAGCCCAGATCCT |
| TGFB3 | Transforming growth factor-β3 | Cell chemotaxis and angiogenesis | F: CAAATTCAAAGGTGTGGACAGT  R: TGAGGGCTGTGTTCCTTCTT |
| TIMP1 | Tissue inhibitor of metalloproteinase 1 | Matrix remodelling | F: CTGCGGATACTTCCACAGGT  R: CAAAACTGCAGGTGGTGATG |
| TIMP1 | Tissue inhibitor of metalloproteinase 1 | Matrix remodelling | F: TCTATGCTGCTGGCTGTGAG  R: GGTCTGTCCACAAGCAGTGA |
| TIMP2 | Tissue inhibitor of metalloproteinase 2 | Matrix remodelling | F: TGATCCCCTGCTACATCTCC  R: GTGCCCGTTGATGTTCTTCT |
| TIMP2 | Tissue inhibitor of metalloproteinase 2 | Matrix remodelling | F: AGAAGAGCCTGAACCACAGG  R: TCCGGAGAGGAGATGTAGCA |
| TIMP3 | Tissue inhibitor of metalloproteinase 3 | Matrix remodelling | F: CTGACAGGCCGTGTCTATGA  CTGGGAGAGGGTGAGCTG |
| TIMP3 | Tissue inhibitor of metalloproteinase 3 | Matrix remodelling | F: GCACACTGGTCTACACCATCA  R: TATACTGCACATGGGGCATC |
| TIMP4 | Tissue inhibitor of metalloproteinase 4 | Matrix remodelling | F: GCTGCCAAATCACCACCT  R: GTCTGTCCAGAGGCACTCG |
| TIMP4 | Tissue inhibitor of metalloproteinase 4 | Matrix remodelling | F: CTCTTGACTGGTCAGATCCTCAG  R: TTCTCCCAGGGCTCAATGT |
| TNF | Tumour necrosis factor-α | Macrophage activation | F: CCCCCAGAAGGAAGAGTTTC  R: CGGGCTTATCTGAGGTTTGA |
| VCAN | Versican | Extracellular matrix molecule | F: GCAGCACACTGCAATATGAGA  R: CACAACGCAGTCTTCTCCAG |
| VCAN | Versican | Extracellular matrix molecule | F: GTGAGCAAGACACGGAGACAT  R: TGGGCGAAGTACTTGTAGCAC |
| VEGF-A | Vascular endothelial growth factor A | Angiogenesis | F: CGAAGGTCTGGAGTGTGTGC  R: TCTCTCCTATGTGCTGGCCT |
| *ACTB* | Actin, beta | Reference gene | F: CTACGTCGCCCTGGACTTC  R: GCAGCTCGTAGCTCTTCTCC |
| *B2M* | Beta-2-microglobulin | Reference gene | F: TGAAGCACGTGACTCTCGAT  R: CTCTGTGATGCCGGTTAGTG |
| *GAPD* | Glyceraldehyde-3-phosphate dehydrogenase | Reference gene | F: ACCCAGAAGACTGTGGATGG  R: AAGCAGGGATGATGTTCTGG |
| *HPRT1* | Hypoxanthine phosphoribosyl transferase I | Reference gene | F: ACACTGGCAAAACAATGCAA  R: TGCAACCTTGACCATCTTTG |
| *RPL13A* | Ribosomal protein L13a | Reference gene | F: ATTGTGGCCAAGCAGGTACT  AATTGCCAGAAATGTTGATGC |
| *PPIA* | Peptidylprolyl isomerase A (cyclophilin A) | Reference gene | F: CAAGACTGAGTGGTTGGATGG  R: TGTCCACAGTCAGCAATGGT |
| *YWHAZ* | Tyrosine 3-monooxygenase/tryptophan 5-monooxygenase | Reference gene | F: GCTGCTGGTGATGATAAGAAGG  R: AGTTAAGGGCCAGACCCAAT |

**Supplementary table 1**: Gene symbol, gene name, gene function and sequences (5’ to 3’) of forward (F) and reverse (R) primers. 56 genes of interest were studied, and 7 reference genes were used. Two sets of primers were designed for genes that had not previously been tested.
