## Supplementary figures and images for "Gene expression profiling for forensic age assessment of porcine skin wounds"

### Supplementary Figure 1

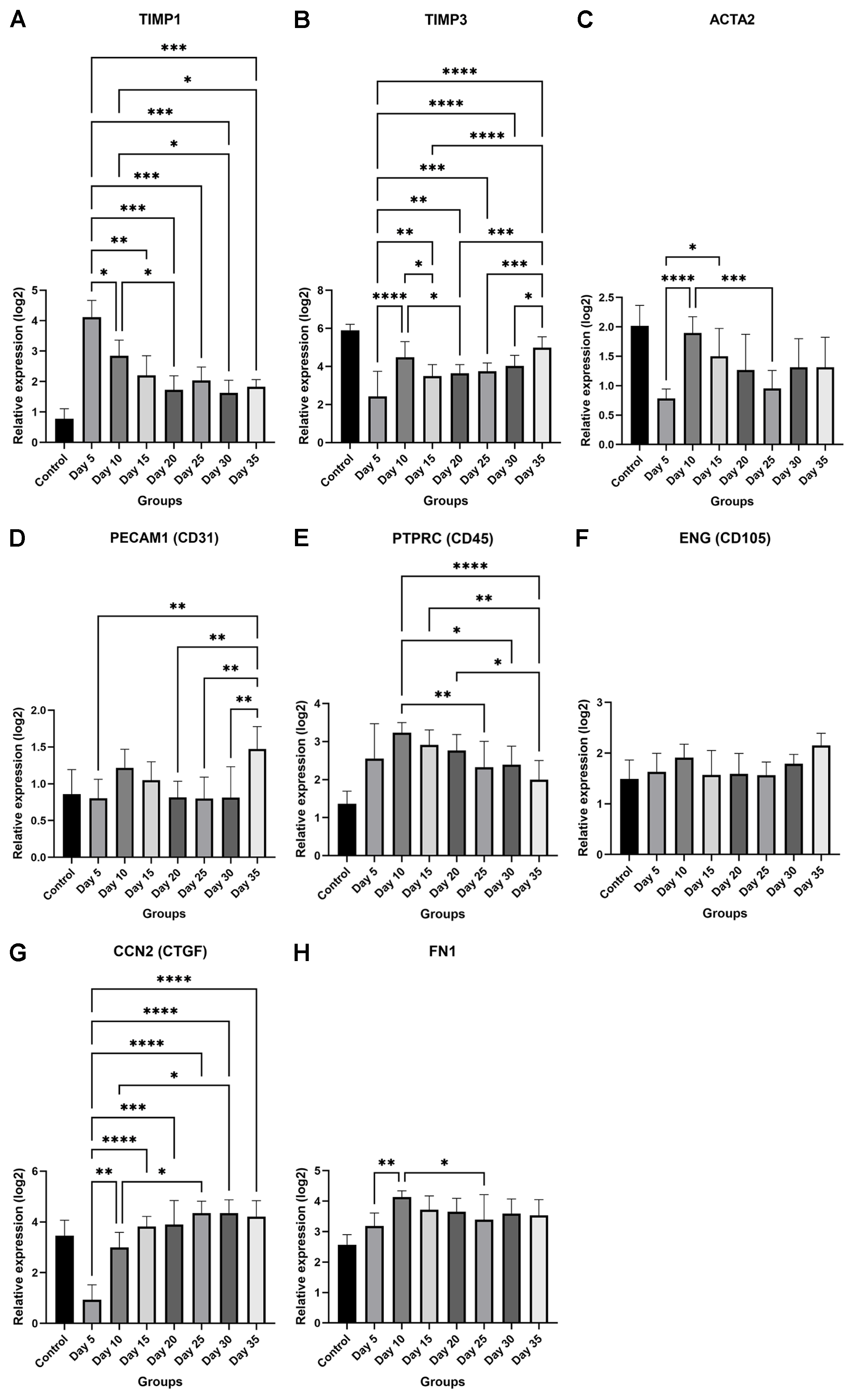
